# Does Infarct Size Influence Gut Barrier Integrity and Bacterial Translocation after Experimental Stroke?

**DOI:** 10.1101/2025.11.13.688219

**Authors:** Cristina Granados-Martinez, Nuria Alfageme-Lopez, Alba Fernandez-Bustelo, Monica A. Amarilla, Manuel Navarro-Oviedo, David Sevillano-Fernandez, Luis Alou-Cervera, Olivia Hurtado, Maria A. Moro, Ignacio Lizasoain, Jesus M. Pradillo

## Abstract

**Background:** Ischemic stroke (IS) triggers brain injury and systemic changes, including gut dysbiosis, intestinal barrier dysfunction (IBD), and bacterial translocation (BT). Although larger infarcts are associated with higher infection risk, it is unclear whether lesion size directly influences gut barrier integrity or bacterial dissemination.

**Methods:** Male C57BL/6 mice underwent permanent middle cerebral artery occlusion (MCAO) proximally or distally to generate large or small infarcts. Seventy-two hours post-ischemia, infarct volume was measured by MRI, and BT was assessed in mesenteric lymph nodes, liver, spleen, and lungs. ZO-1 and MMP9 expression were used to evaluate intestinal barrier integrity. Peripheral and central inflammation were assessed by flow cytometry and immunofluorescence.

**Results:** Proximal MCAO produced larger infarcts than distal MCAO. The proportion of animals exhibiting BT was lower in distal MCAO, but this difference was not statistically significant. ZO-1 expression did not differ between groups, while MMP9 was increased only in animals with BT, independent of infarct size. BT was associated with more pronounced lymphopenia and enhanced microglial activation and T-cell infiltration in the brain. The composition of translocated bacteria was similar across groups.

**Conclusions:** Infarct size alone does not determine IBD or BT, although BT is linked to intestinal and cerebral inflammation. This study evaluates for the first time the effect of lesion magnitude on IBD and BT, highlighting the complex interplay between cerebral injury and gut–systemic interactions.

## Introduction

Stroke is a leading cause of death and long-term disability worldwide, representing a major global health burden^1^. Ischemic stroke (IS), which accounts for the majority of cases, not only produces brain tissue damage and disruption of the blood–brain barrier (BBB), but also triggers systemic alterations that affect peripheral organs^1,2^. Among these systemic consequences, profound changes in the gut microbiota, increased intestinal permeability, and bacterial translocation (BT) to extra-intestinal tissues have been increasingly recognized^1-3^. Experimental models of middle cerebral artery occlusion (MCAO) have shown that stroke induces gut dysbiosis and alters epithelial tight junction proteins such as claudins and ZO-1, leading to intestinal barrier disruption and promoting bacterial translocation (BT) to mesenteric lymph nodes (MLN), liver, spleen, and lungs^2-5^. This BT has been associated with worsened stroke outcomes and identified as a contributing factor to the development of post-stroke infections^6^.

Clinically, larger infarct volumes are associated with worse functional outcomes, stronger systemic inflammatory responses, and a higher incidence of infections after stroke^7,8^. Infarct size has also been implicated in BBB disruption and post-stroke immunodepression, which may facilitate bacterial invasion^7,8^. Despite this evidence, no experimental study has directly evaluated whether the initial infarct size determines the magnitude of intestinal barrier disruption (IBD) or BT. Understanding this relationship is crucial to determine whether a threshold of cerebral injury triggers gut barrier failure and systemic infection, potentially amplifying peripheral and central inflammation and influencing infarct progression. Such insights could open new therapeutic avenues targeting the gut barrier or microbiota, particularly in patients with extensive ischemic lesions^9^.

## Methods

### Animals

Experiments were conducted in male C57BL/6 wild-type mice (C57BL/6J Ola Hsd; Envigo, Spain) aged 8–10 weeks and weighing 20–30 g. Animals were housed under controlled conditions (22 °C, 35% humidity, 12 h light/dark cycle) with free access to food and water. All procedures complied with the European Communities Council Directive (86/609/EEC) and were approved by the Animal Welfare Ethics Committee of the Complutense University (PROEX No. 219.3/24). Experiments followed ARRIVE guidelines, and animal numbers were minimized using prior experience and statistical tools (http://www.biomath.info). Overall mortality was below 7%.

### Experimental design and induction of ischemia

Animals were randomly assigned to *naïve* or stroke (MCAO) groups, and all analyses were performed in a blinded manner. Focal IS was induced by permanent electrocoagulation of the left middle cerebral artery (MCA) as previously described^10^. To generate different infarct volumes, the MCA was occluded either proximally, before bifurcation of the frontal and parietal branches to produce large infarcts, or distally, by coagulation of the parietal branch to yield smaller lesions^11^.

Based on prior evidence indicating that BT occurs 48–72h after stroke^4,9^, all animals were euthanized, and analyses performed 72h post-ischemia. Following microbiological assessment, animals were classified as exhibiting BT or no BT (NBT).

### Infarct volume and neurological evaluation

At 72h, infarct volume was quantified using T2-weighted MRI (Icon 1T-MRI; Bruker BioSpin GmbH, Germany). Anesthetized mice underwent 3D RARE imaging (TR = 2200 ms; TE = 75 ms; FOV = 34 × 31 × 12 mm; voxel size = 0.25 × 0.25 × 0.50 mm; total time ≈ 10 min). Infarct size was expressed as the percentage of the ischemic hemisphere (%IH) as described previously^12^.

### Bacterial translocation analysis

At 72h, MLN, liver, spleen, and lungs were aseptically harvested, weighed, and homogenized in sterile saline. Homogenates were plated on Columbia and MacConkey agar for aerobic culture (35 °C, 48h) and on Brucella agar for anaerobic growth (35 °C, 96h) (Becton Dickinson, NJ, USA). Colony-forming units (CFU) were quantified and expressed as log_10_ CFU/g tissue (detection limit = 1.3 log_10_ CFU/g). Distinct colonies were identified by morphology, subculture, and API biochemical tests, following the *International Code of Nomenclature of Prokaryotes*^13^.

### Peripheral and central inflammation

Peripheral inflammation was assessed by flow cytometry of bone marrow (BM) and blood leukocytes stained with specific antibody panels and analyzed with FlowJo v10 (BD Biosciences). Central inflammation was evaluated by immunofluorescence on brain sections stained for microglia (IBA-1), T cells (CD3), and endothelial cells (CD31). Images were acquired by confocal microscopy and quantified using ImageJ.

### Statistical analysis

Data are presented as mean ± SD or median [IQR] as appropriate. Normality was assessed using the Kolmogorov–Smirnov test. For two-group comparisons, Student’s t-test or Mann–Whitney U test was applied. For multiple groups, Kruskal– Wallis with Dunn’s post hoc test was used. Shannon and Simpson indices were used to assess BT diversity and dominance across organs. Analyses were performed using GraphPad Prism 8.0.1 and R (v4.4.2); p < 0.05 was considered statistically significant.

## Results

### Infarct Volume, Bacterial Translocation, and Intestinal Barrier Integrity

Animals subjected to distal MCAO developed significantly smaller infarcts than those with proximal occlusion, while both models exhibited comparable neurological and behavioral impairments 72h after surgery, confirming the functional impact of ischemia (Fig. 1A).

**Figure 1.**
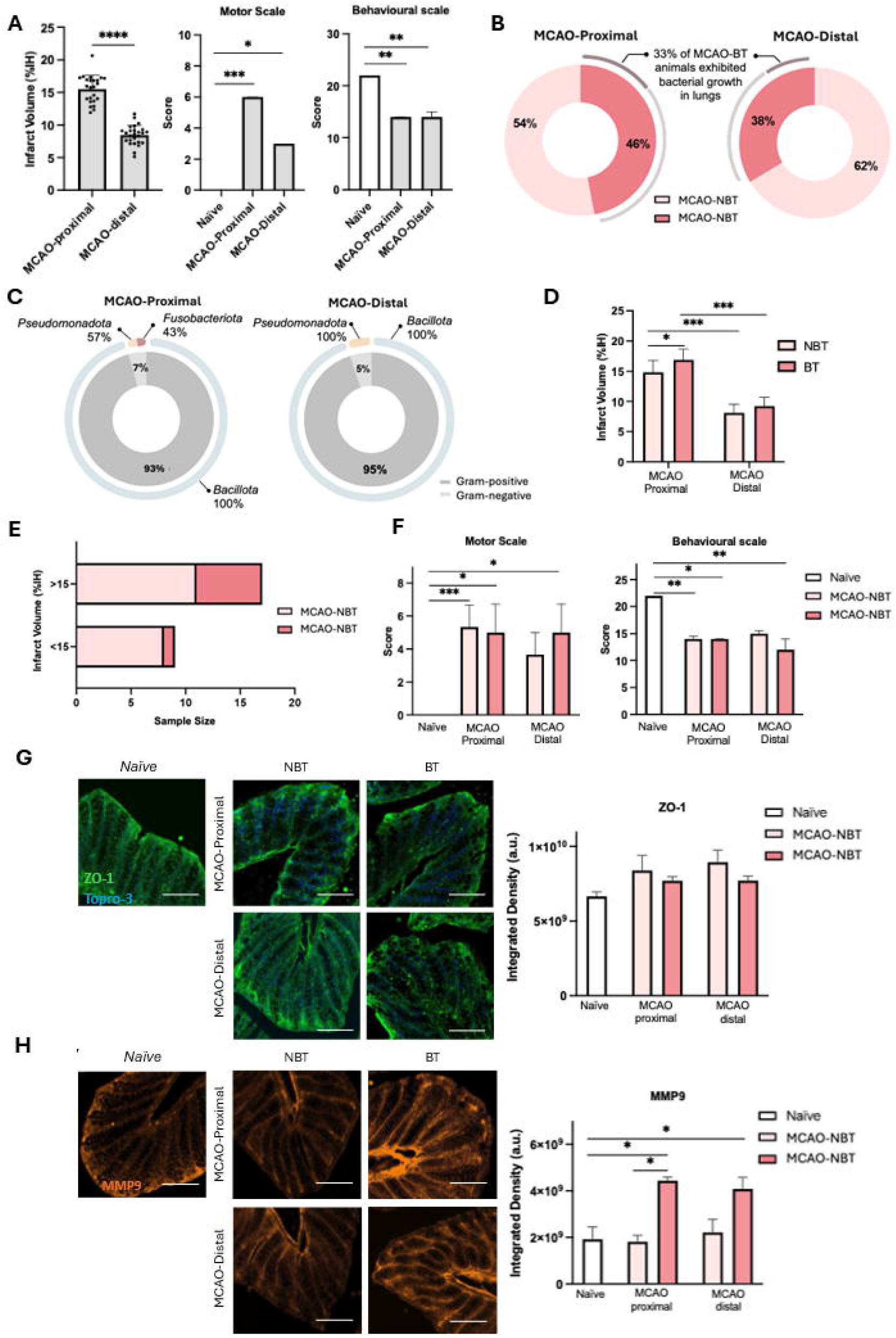
Infarct volume, stroke outcome, bacterial translocation and intestinal barrier damage after experimental ischemic stroke. (A) Infarct volume (expressed as the percentage of the ipsilateral hemisphere, %IH) at 72h post-surgery in MCAO-proximal and MCAO-distal animals (n = 24–26). Results are shown as mean ± SD; Student’s *t*-test, *p* < 0.01. Motor and behavioural characterization in naïve (n = 6) and at 72h in MCAO-proximal and MCAO-distal (n = 24–26) animals. Results are expressed as median (IQR). Statistical analysis was performed using the Kruskal–Wallis test followed by Dunn’s post hoc multiple comparisons test; *p* < 0.05, \**p* < 0.01, \*\**p* < 0.001, \*\*\**p* < 0.0001. (B) Percentage of animals showing bacterial translocation (BT). (C) Study of the relative abundance of Gram-positive and Gram-negative bacteria, and the different bacterial phyla isolated from the organs analyzed 72 hours after experimental ischemic stroke in MCAO-proximal and MCAO-distal animals with bacterial translocation. (D) Infarct volume (expressed as %IH) at 72h post-surgery in MCAO-proximal and MCAO-distal groups in NBT (n = 16–18) and BT (n = 8) animals. Results are shown as mean ± SD; two-way ANOVA followed by Bonferroni post hoc multiple comparisons test; *p* < 0.05, \*\**p* < 0.001. (E) Number of animals developing BT at 72h post-stroke as a function of infarct volume greater or smaller than 15% of the ipsilateral hemisphere. (F) Motor and behavioral characterization in naïve (n=4) and at 72h in MCAO-proximal and MCAO-distal groups in NBT (n = 16-18) and BT (n = 8) animals. Results are expressed as median (IQR). Statistical analysis was performed using the Kruskal–Wallis test followed by Dunn’s post hoc multiple comparisons test; *p < 0.05, **p < 0.01, ***p < 0.001. (G) Integrated density (a.u.) of ZO-1 levels in the colon quantified in naïve (n = 4) and at 72h in MCAO-proximal and MCAO-distal groups in NBT (n = 16–18) and BT (n = 8) animals. ZO-1 (green), Topro-3 (blue). Scale bar, 50 μm. Results are shown as median (IQR); Kruskal–Wallis test followed by Dunn’s post hoc multiple comparisons test. (H) Integrated density (a.u.) of MMP9 levels in the colon quantified in naïve (n = 4) and at 72h in MCAO-proximal and MCAO-distal groups in NBT (n = 16–18) and BT (n = 8) animals. MMP9 (red). Scale bar, 50 μm. Results are shown as median (IQR); Kruskal–Wallis test followed by Dunn’s post hoc multiple comparisons test; *p* < 0.05.

The incidence of BT tended to be lower in the distal MCAO group, although this difference did not reach statistical significance (Fig. 1B). Bacterial dissemination to the lungs occurred with similar frequency in both models. The taxonomic composition of translocated bacteria was also comparable, with no significant differences in Gram-type distribution (Fig. 1C). Within Gram-positive taxa, *Bacillota* was consistently detected in both models. Among Gram-negative bacteria, *Pseudomonadota* and *Fusobacteriota* were identified in proximal MCAO, whereas only *Pseudomonadota* was detected in distal MCAO.

When infarct size was stratified by BT status, proximal MCAO animals with BT displayed significantly larger lesions than non-translocating counterparts, whereas no effect was observed in the distal model (Fig. 1D). To further explore this association, proximal MCAO animals were categorized by infarct volume. Although not statistically significant, BT occurrence tended to be higher in mice with infarcts exceeding 15% of the hemisphere compared with smaller lesions (Fig. 1E). Importantly, BT did not influence neurological or behavioral performance in either model (Fig. 1F).

Regarding intestinal barrier integrity, ZO-1 expression remained unchanged across experimental conditions (Fig. 1G). In contrast, MMP9 expression was markedly elevated in ischemic animals exhibiting BT compared with both non-translocating and naïve controls, irrespective of infarct size or occlusion site (Fig. 1H).

### Peripheral and Central Inflammatory Responses According to Infarct Volume and Bacterial Translocation

Flow cytometric analysis of BM revealed no significant differences in myeloid or lymphoid populations between proximal and distal MCAO groups, regardless of BT status (Fig. 2A). In contrast, peripheral blood analysis showed a marked reduction in total lymphocytes after stroke, more pronounced in animals with BT, accompanied by a trend toward increased monocyte counts (Fig. 2B–C).

**Figure 2.**
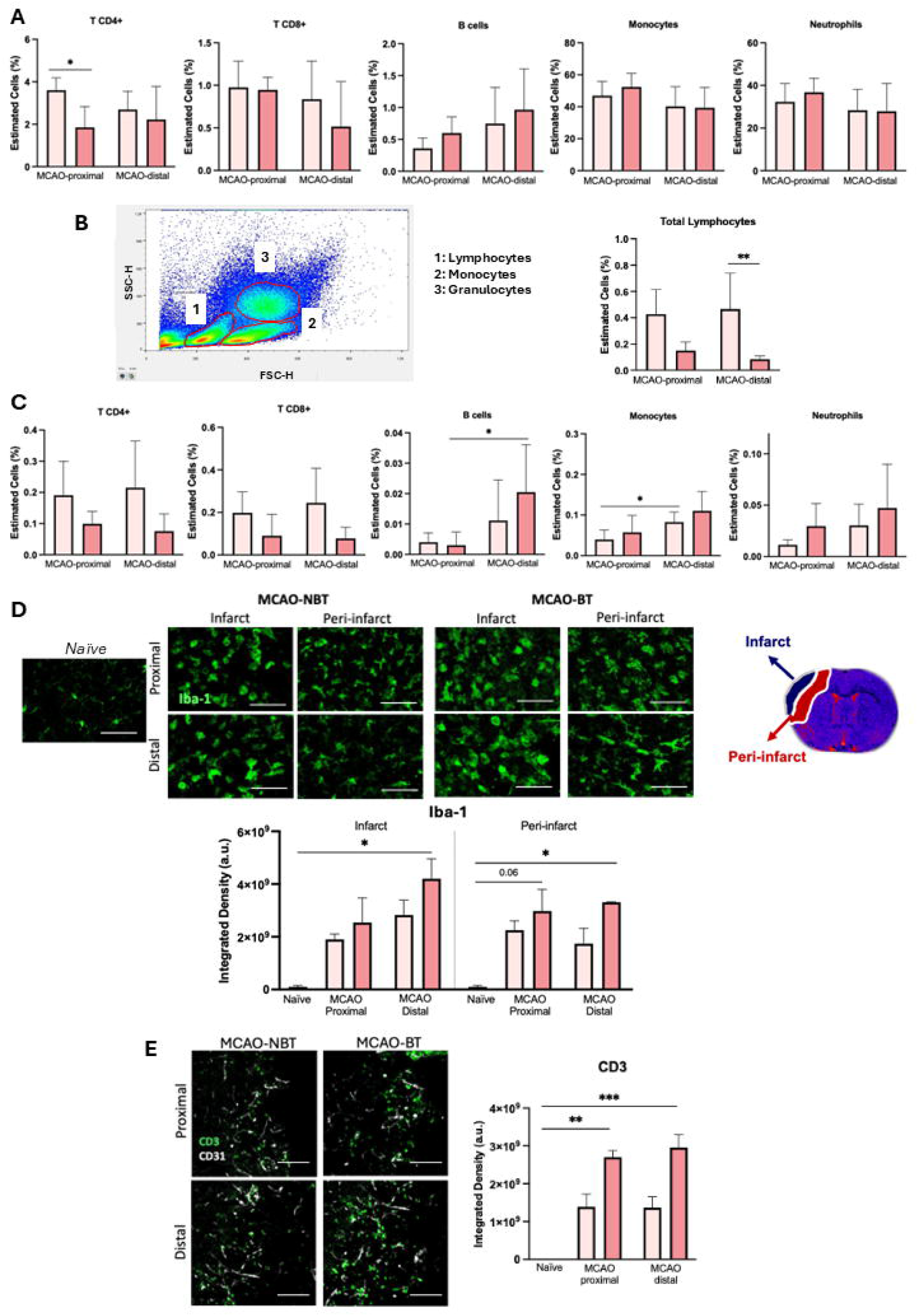
Impact of bacterial translocation on the peripheral and central inflammatory response at 72h after experimental ischemic stroke. Immune cell levels assessed by flow cytometry in (A) bone marrow and (B, C) peripheral blood at 72h post-surgery in MCAO-proximal and MCAO-distal groups in NBT (n = 16–18) and BT (n = 8) animals. Results are shown as median (IQR); Kruskal–Wallis test followed by Dunn’s post hoc multiple comparisons test, *p < 0.05, **p < 0.01. (D) Integrated density of Iba-1 in the brain, quantified in naïve (n=4) and at 72h after ischemic stroke in MCAO-proximal and MCAO-distal groups in NBT (n = 16–18) and BT (n = 8) animals. Results are shown as median (IQR); Kruskal–Wallis test followed by Dunn’s post hoc multiple comparisons test, *p < 0.05. (E) Integrated density of CD3^+^ lymphocyte infiltration quantified in naïve (n=4) and at 72 h after ischemic stroke in MCAO-proximal and MCAO-distal groups in NBT (n = 16–18) and BT (n = 8) animals. Results are shown as median (IQR); Kruskal–Wallis test followed by Dunn’s post hoc multiple comparisons test, **p < 0.01, ***p < 0,001.

Within the brain, microglial activation tended to increase in the infarct core of MCAO-proximal animals and was significantly elevated in MCAO-distal animals compared with naïve controls, with a similar pattern in the periinfarct region (Fig. 2D). Consistently, T-cell (CD3^+^) infiltration into the infarcted area was greater in MCAO-proximal than MCAO-distal animals (Fig. 2E).

Overall, our results indicate that infarct size shapes both systemic and central inflammatory responses, while IBD and BT are linked to amplified neuroimmune activation after IS.

## Discussion

In this study, we explored the relationship between cerebral infarct size, IBD and BT following IS. Consistent with previous work from our laboratory, animals subjected to distal MCAO exhibited smaller infarct volumes at 72h post-ischemia, similar to the reduced lesion size reported at 24h in this model^11^. Although proximal MCAO produced larger infarcts, the proportion of animals exhibiting BT was not significantly different between groups, with only a non-significant trend toward lower BT in animals with smaller lesions. These findings suggest that infarct size alone may not be the primary determinant of gut barrier compromise or bacterial dissemination, consistent with prior evidence that post-stroke gut dysbiosis and barrier dysfunction involve multiple factors, including autonomic dysregulation, intestinal motility, and systemic inflammation^2,5,6^.

Analysis of IBD showed no differences in ZO-1 expression across groups with different lesion sizes. In contrast, MMP9 levels, a marker of intestinal inflammation and barrier degradation, were elevated specifically in animals with BT, regardless of infarct volume. This indicates that post-stroke intestinal inflammation and barrier dysfunction may be more closely associated with BT itself than with the extent of cerebral injury, in line with previous studies implicating MMP9 in facilitating microbial dissemination^4,5^.

Peripheral lymphopenia was more pronounced in animals with BT, and central neuroinflammation, including microglial activation and T-cell infiltration, was enhanced in these animals. Notably, microglial activation tended to increase in the infarct core of MCAO-proximal animals and was significantly elevated in MCAO-distal animals, with a similar pattern observed in the periinfarct region, whereas T-cell infiltration was higher in proximal MCAO animals. These findings support the concept of bidirectional gut–brain communication, in which BT may amplify neuroimmune responses^2,3^.

Overall, our results indicate that infarct size does not directly dictate IBD or BT, although larger lesions may predispose to BT in some cases. Importantly, this study evaluates for the first time the effect of lesion magnitude on IBD and BT, providing a framework for future investigations into gut–brain interactions after stroke. Therapeutic strategies targeting intestinal inflammation or microbiota modulation may benefit patients prone to post-stroke infections, independently of infarct size^9^.

## Acknowledgments

This research was funded by grants from the Spanish Ministry of Science and Innovation (PID2020-117765RB-I00; Dr Pradillo; PID2022-140616OB-I00, Dr Moro), Leducq Trans-Atlantic Network of Excellence (TNE-21CVD04; Drs Lo, Moro, Lizasoain), Instituto de Salud Carlos III, the European Development Regional Fund, RICORS-ICTUS (RD21/0006/0001), and the FORTALECE program (FORT23/00023; Dr Lizasoain).

## References

1. Stanley D, Moore RJ, Wong CHY. An insight into intestinal mucosal immunity and microbiota disruption after stroke. J Neuroinflammation. 2018;15(1):201.

2. Singh V, Roth S, Llovera G, et al. Microbiota dysbiosis controls the neuroinflammatory response after stroke. J Neurosci. 2016;36(28):7428– 40.

3. Benakis C, Brea D, Caballero S, et al. Commensal microbiota affects ischemic stroke outcome by regulating intestinal γδ T cells. Nat Med. 2016;22(5):516–23.

4. Crapser J, Ritzel R, Verma R, et al. Ischemic stroke induces gut permeability and enhances bacterial translocation leading to sepsis in aged mice. Aging (Albany NY). 2016;8(5):1049–63.

5. Houlden A, Goldrick M, Brough D, et al. Brain injury induces specific changes in the caecal microbiota of mice via altered autonomic activity and mucoprotein production. Brain Behav Immun. 2016;57:10–20.

6. Chamorro Á, Urra X, Planas AM. Infection after acute ischemic stroke: A manifestation of brain-induced immunodepression. Stroke. 2007;38(3):1097–103.

7. Haeusler KG, Schmidt WU, Föhring F, et al. Inflammation and infection after ischemic stroke: Correlation with infarct volume and outcome. Stroke. 2008;39(10):2824–30.

8. Westendorp WF, Nederkoorn PJ, Vermeij JD, et al. Post-stroke infection: A systematic review and meta-analysis. BMC Neurol. 2011;11:110.

9. Spychala MS, Venna VR, Jandzinski M, et al. Age-related changes in the gut microbiota influence systemic inflammation and stroke outcome. Ann Neurol. 2018;84(1):23–36.

10. Caso JR, Pradillo JM, Hurtado O, et al. Toll-like receptor 4 is involved in brain damage and inflammation after experimental stroke. Circulation. 2007;115(12):1599–608.

11. Moraga A, Pradillo JM, Cuartero MI, et al. Toll-like receptor 4 modulates cell death and inflammation after experimental stroke. FASEB J. 2014;28(11):4710–18.

12. Hernández-Jiménez M, Gómez-Sánchez JC, Falcón D, et al. Silencing of the mitochondrial protein mortalin protects against ischemic brain injury. Stroke. 2013;44(8):2240–48.

13. Oren A, Garrity GM. Valid publication of new names and new combinations effectively published outside the IJSEM. Int J Syst Evol Microbiol. 2021;71(11):e005095.

14. Iadecola C, Anrather J. The immunology of stroke: From mechanisms to translation. Nat Med. 2011;17(7):796–808.

15. Meisel C, Schwab JM, Prass K, et al. Central nervous system injury-induced immune deficiency syndrome. Nat Rev Neurosci. 2005;6(10):775–86.

